# Non-apoptotic role of EGL-1 in exopher production and neuronal health in *Caenorhabditis elegans*

**DOI:** 10.1101/2024.04.19.590348

**Authors:** Zheng Wu, Eric A. Cardona, Jonathan T. Pierce

## Abstract

While traditionally studied for their pro-apoptotic functions, recent research suggests BH3-only proteins also have non-apoptotic roles. Here, we find that EGL-1, the BH3-only protein in *Caenorhabditis elegans*, promotes the cell-autonomous production of exophers in adult neurons. Exophers are large, micron-scale vesicles that are ejected from the cell and contain cellular components such as mitochondria. EGL-1 facilitates exopher production potentially through regulation of mitochondrial dynamics. Moreover, an endogenous, low level of EGL-1 expression appears to benefit dendritic health. Our findings provide insights into the mechanistic role of BH3-only protein in mitochondrial dynamics, downstream exopher production, and ultimately neuronal health.

**Significance statement:** BH3-only proteins were known for their function in inducing cell death. Their presence in healthy adult neurons, however, suggests additional roles. Our study focused on the BH3-only protein EGL-1 in the nematode *Caenorhabditis elegans*, where its apoptotic role was discovered. We reveal a new role in cell-autonomously promoting exopher production – a process where neurons extrude large vesicles containing potentially harmful cell contents. EGL-1 appears to promote this by regulating mitochondrial dynamics. We also report that low levels of EGL-1 benefit neuronal health and function. These findings expand our understanding of BH3-only proteins, mitochondrial dynamics, and exopher production in neurons and provide insights for neurodegenerative diseases.

## Introduction

Apoptosis is important for both developmental processes and homeostatic mechanisms in metazoans. The apoptosis pathway involves Bcl-2 Homology (BH) 3-only proteins, which are conserved and function by inhibiting BCL-2 proteins, releasing components necessary for caspase activation^[1]^. In *C. elegans*, EGL-1 is the BH3-only protein that serves as the primary initiator of apoptosis^[2,3]^. While the BH3-only proteins have been studied in their pro-apoptotic roles, recent studies suggest they also have non-apoptotic functions^[4]^. Mammalian studies have reported the endogenous expression of BH3-only proteins in the nervous system at low basal levels^[5,6,7]^. While these low basal expression levels are insufficient to induce apoptosis, their specific functions remain largely unknown.

In a previous study, our lab discovered that *egl-1* is highly expressed in URX neurons in adult *C. elegans* without causing apoptosis, and that *egl-1* expression is cell-autonomously dependent on neuronal activity^[8]^. Here we report that *egl-1* is required cell-autonomously for efficient production of exophers in URX neurons. Exophers are large (>2 μm diameter), membrane-bound vesicles that are extruded from neurons and have been found to contain protein aggregates and organelles such as mitochondria^[9]^.

The highly dynamic nature of exophergenesis necessitates *in vivo* observations to study the budding, ejection, release, phagocytosis, and degradation of the exopher^[10**Error! Bookmark not defined**.]^. Large extracellular vesicles observed in other model organisms are often recognized as apoptotic bodies or artifactual cell debris from the tissue fixation process in immunohistochemistry, making it hard to determine their identity and origin^[11]^. The transparent body and limited number of somatic cells in *C. elegans* make it a suitable model organism for visualizing the study of exophers *in vivo*.

Here, we studied exophergenesis in URX and the well-established ALMR touch neuron exopher model using *in vivo* microscopy, gene knock-down and rescue, and epistasis analyses. We found that *egl-1* promotes exopher production cell-autonomously in both classes of neurons, potentially by promoting mitochondrial fission. Consistent with previous exopher studies^[9]^, we found evidence supporting that exopher production seems to be beneficial for neuronal health. Specifically, inhibition of basal non-apoptotic neuronal *egl-1* expression appeared detrimental to URX dendritic health. Our findings connecting the *egl-1*-regulated mitochondrial dynamics and exopher production provide novel insights into the non-apoptotic functions of BH3-only proteins and exopher production in neurons.

## Results

Exophers have been identified in multiple neuron types since their discovery in *C. elegans*^[9]^. These include many of the touch neurons and additional sensory neurons including CEP and ASER in worm^[9]^. While studying the oxygen-sensing neuron pair URX, we found that it also produced exophers under normal physiological conditions (Figure 1A). URX exophers were consistently large (average area 5.0 μm^2^) and roughly one-fourth the size of the URX soma (average area 16.4 μm^2^; Supplemental Figure 1). URX exophers contained mitochondria, as identified by mitoGFP, like previously reported ALMR exophers (Figure 1C). When analyzing hundreds of URX exophers, we found that they are often produced from the posterior and dorsal side of the soma, which is on the side opposite from where the URX dendrite and axon originate (Figure 1A and 1E). While the well-studied ALMR exophers have a generally spherical shape^[9]^, URX exophers are morphologically more variable (Figure 1B). In our observation, while the URX soma remains relatively static, URX exophers display highly dynamic morphologies when the animals are allowed to crawl slowly during imaging.

**Figure 1.**
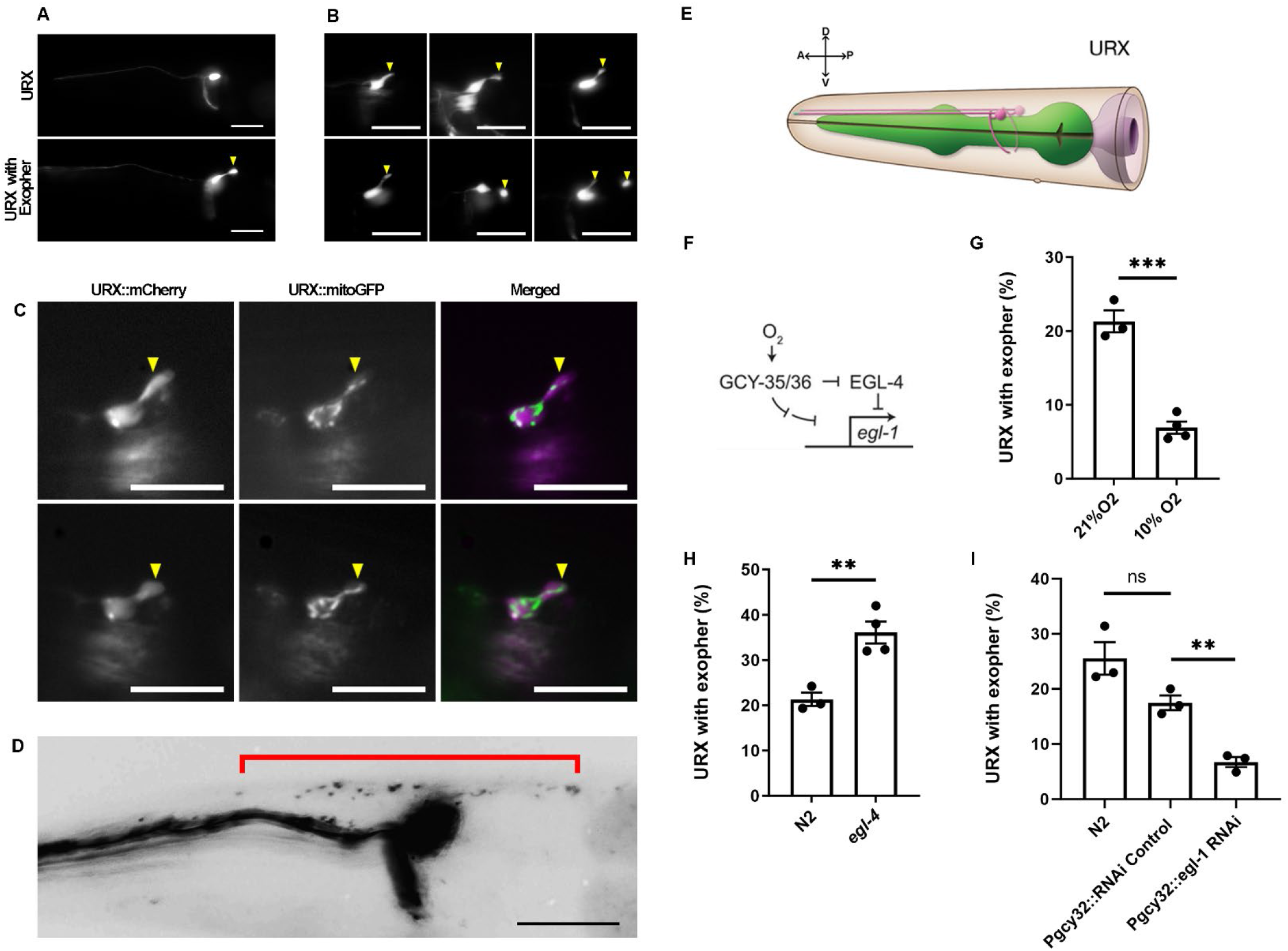
*egl-1* facilitates exopher production in URX neurons. A. Representative images of a URX neuron (top) and a URX neuron with an exopher from a different animal (bottom). URX is visualized with *Ex[Pgcy32::GFP]* reporter. B. URX exophers have highly variable morphologies. URX is visualized with *Ex[Pgcy32::GFP]* reporter. C. URX exophers contain mitoGFP-labeled mitochondria. Top: Mitochondria in the exopher appear discontinuous from the somatic mitochondria. Bottom: Mitochondria appear contiguous in the exopher and soma. Left panels show URX with mCherry reporter, middle panels show mitoGFP, right panels are merged images. Strain is *Is[Pegl-1::mCherry];Is[Pegl-1::mitoGFP]*. D. URX-released exopher appears to be phagocytosed by the surrounding hypodermis and undergoes tubulation and vesiculation. URX and its released exopher contents are visualized with *Ex[Pgcy32::mCherry]* reporter. E. Illustration of URX neuron from WormAtlas, all URX images in this paper are aligned within the same direction. F.*egl-1* expression in URX is dependent on O_2_ and Ca^2+^ and suppressed by *egl-4*. G. Wild-type N2 animals are grown in 21% or 10% O_2_ environment and are scored for URX exophers. URX displays fewer exophers when grown in lower O_2_ environment, where URX has lower Ca^2+^ activity and lower *egl-1* expression. URX is labeled with *Is[Pgcy32::GFP]*. Dots represent groups with N = 247, 417 worms total for 21% and 10% O_2_ environment, respectively. H. URX exophers are scored in *egl-4(n477)* and compared to N2. URX has more exophers in *egl-4* loss-of-function mutants, in which URX has higher *egl-1* expression. URX is labeled with *Is[Pgcy32::GFP]*. N = 247, 215 worms total for wild-type and mutant respectively. I. URX exophers are scored in N2, *Ex[Pgcy32::GFP, Pgcy32::mCherry RNAi]* and *Ex[Pgcy32::GFP, Pgcy32::egl-1 RNAi]*. URX-specific knockdown of *egl-1* reduces exopher production, while the control strain with URX-specific knockdown of mCherry is not significantly different from the N2. N = 185, 189 and 191 worms total for wild-type, RNAi control and *egl-1 RNAi* respectively. All animals are adult day 1. Scale bars: 20 μm.

A recent study showed that ALMR exophers are stimulated and engulfed by the surrounding hypodermis via a phagocytotic process^[12]^. The engulfed exopher degrades to yield hallmarks including tubulation and vesiculation to form “starry night” vesicles throughout the tissue^[12]^. Utilizing a mCherry-labeled URX strain we identified a similar “starry night” pattern in the hypodermis near URX (Figure 1D), suggesting a comparable exopher production and degradation mechanism for URX. This “starry night” pattern is exclusive to neurons labeled with mCherry, contrasting those labeled with GFP. Considering GFP is more susceptible to acidic environments compared to mCherry ^[13]^, this observation suggests lysosomic degradation of exophers by the surrounding tissue. We propose that the high variability in URX exopher morphology likely results from the different spatial relationship between the neuron soma and the hypodermis; the ALM touch neuron is embedded in the hypodermis, whereas the URX neuron is located in the pseudo-coelomic cavity between the dorsal hypodermis and the pharynx^[14]^.

Taken together, our observations show that URX neurons produce exophers, and the phagocytotic and degradation mechanisms of exophers by the adjacent hypodermis may be shared in URX.

### EGL-1 cell-autonomously required for efficient exopher production

URX neurons respond to ambient oxygen level tonically^[15]^. To study the regulation of exopher production in these oxygen-sensing neurons, we raised the animals in environments with different oxygen levels to control URX activity and scored exophers in day 1 adults. We found that when raised in a low (10%) oxygen environment, where URX has low activity, the exopher rate was reduced significantly (Figure 1G). This indicates exopher production is likely activity-dependent in the URX neuron. In our previous study with the URX neuron^[8]^, we found that the apoptosis gene *egl-1* is expressed in URX neurons in healthy adult animals in an activity-dependent manner, and the cGMP-dependent kinase EGL-4 suppresses *egl-1* expression in URX at room oxygen (Figure 1F). Here, we found that URX exopher rate is significantly increased in the *egl-4* mutant (Figure 1H). These results suggest a correlation between *egl-1* expression level and exopher rate in URX.

Does exopher production depend on *egl-1*? This question is hard to address with *egl-1* loss-of-function mutants because, due to their deficiency in apoptosis, they have mis-localized and extra “undead” URX cell bodies that make identification of the original URX neuron and its exophers ambiguous. To surmount this problem, we knocked down *egl-1* cell-specifically in URX via RNAi using a cell-specific (URX, AQR and PQR) promoter^[16]^. Selective knock down of *egl-1* appeared effective because it prevented programmed cell death in the PQR lineage as evident by an extra “undead” PQR neuron in the transgenic animals (Supplemental Figure 2). Due to lineage differences, the knockdown of *egl-1* did not induce extra “undead” URX cells, enabling us to study exophers from URX unambiguously. We found that cell-specific knockdown of *egl-1* significantly decreased the exopher rate in URX compared to a control strain that expressed mCherry dsRNA in the same cells (Figure 1I). This implies a cell-autonomous role of *egl-1* in neuronal exopher production in the URX neuron.

Having identified the involvement of *egl-1* in producing exophers in the URX neuron, we next tested role for apoptotic signaling pathway in exopher production. We started with the touch neuron ALMR, where exophers were discovered^[9]^. In addition to the apoptotic trigger, *egl-1*, we also considered the downstream apoptosis pathway, which includes the Bcl-2 gene *ced-9* and the caspase *ced-3*^[17,18]^. We used a strain with GFP-labeled touch neurons to study the basal exopher rate (Figure 2A). Mutants defective in apoptosis do not have extra “undead” ALMR neurons, so scoring the ALMR exopher rate was unambiguous in these mutants: *egl-1(lf), ced-9(gf)* and *ced-3(lf)*. ALMR exopher rate has been reported to peak at day 2 of adulthood, correlating with the number of fertilized eggs, which are believed to enhance production due to mechanical jostling within the body of the gravid animal^[9,19]^. Apoptosis mutants produce normal fertilized eggs but have slightly smaller brood size and slower growth rate. To ensure the age synchronization and consistency with our URX exopher analysis, we used day 1 adults for scoring ALMR exophers. We found that the ALMR exopher rate is significantly decreased in the *egl-1* loss-of-function and *ced-9* gain-of-function mutants, but not in the *ced-3* mutant (Figure 2B). These results suggest that the BH3-only protein EGL-1 and the BCL-2 homolog CED-9, but not the downstream caspase CED-3, are involved in exopher production.

**Figure 2.**
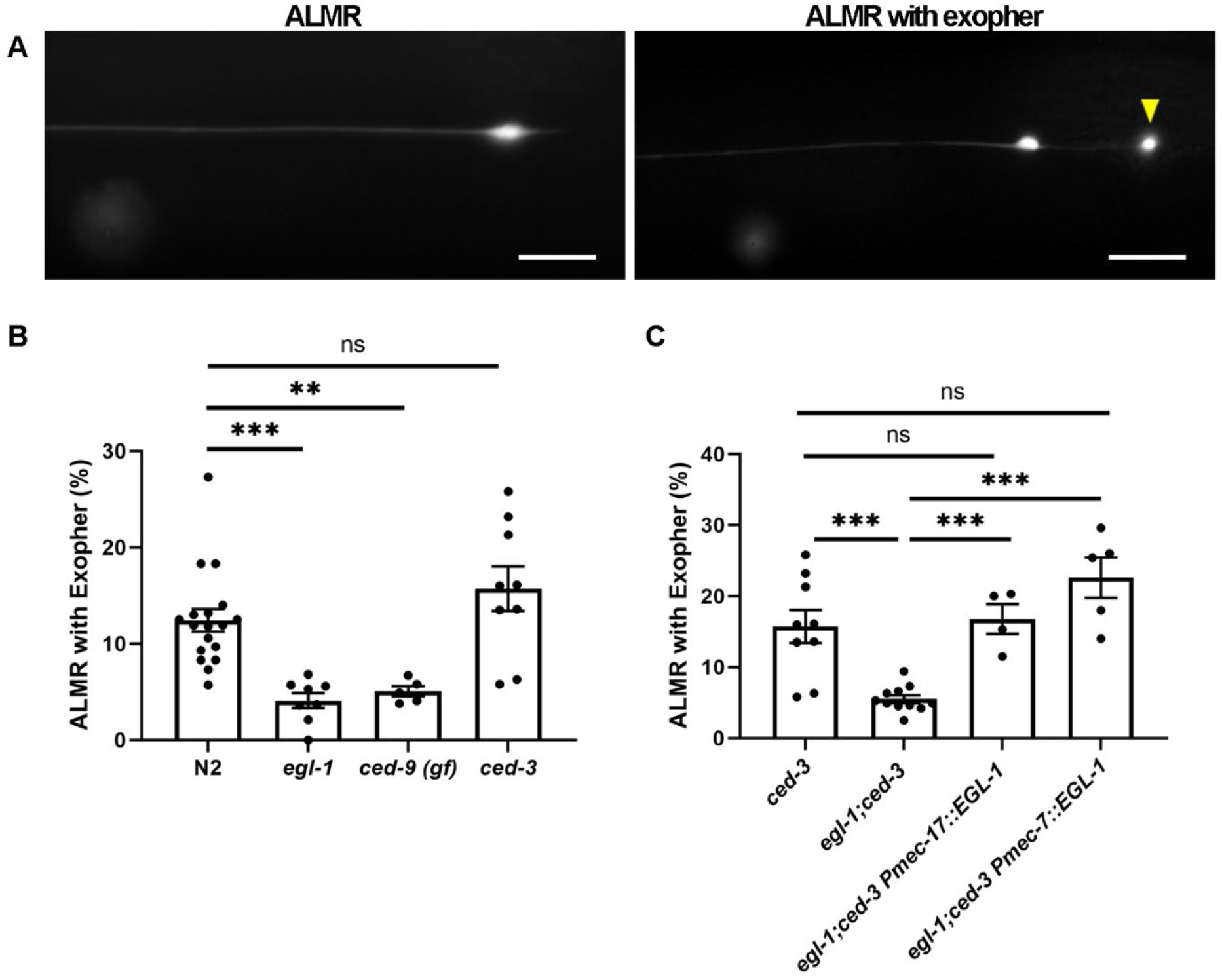
*egl-1* is required in ALMR neuron for efficient exopher production. A. Representative image of ALMR neuron (left) and ALMR neuron with an exopher (right). B. ALMR exophers are scored in wild-type N2, *egl-1(n1084n3082), ced-9 gf(n1950)* and *ced-3(ok2734)* animals. Both *egl-1* loss-of-function and *ced-9* gain-of-function mutants have abnormally fewer exophers, while *ced-3* loss-of-function has no effect on exopher production. Dots represent groups with N = 1082, 708, 317 and 674 worms total for wild-type and each mutant by order, respectively. C. ALMR exophers are scored in *egl-1(n1084n3082)*;*ced-3(ok2734)*, Ex[Pmec-17::EGL-1]; *egl-1(n1084n3082)*; *ced-3(ok2734)* and compared to *ced-3(ok2734)*. Touch-neuron-specific expression of *egl-1* rescued the decreased exopher production caused by *egl-1* loss-of-function mutation. N = 674, 671, 239 and 260 for control, mutant, and rescues respectively. All animals are adult day 1. Touch neurons are labeled in all strains above with integrated GFP reporter *Is[Pmec-17::GFP]*. Scale bars: 20 μm.

Whereas URX neurons express *egl-1* transcript at high levels as determined by smFISH and transcriptional reporters, the ALMR neuron lacks a detectable level of *egl-1* expression as determined by reporter and by a recent single-cell transcriptomic study^[20]^. Genetic evidence, however, importantly suggests the presence of a low but functional level of EGL-1 protein in the ALM neurons. This study reported that the *egl-1* mutation suppressed the ALM neuron death induced by CED-3 overexpression ^[21]^. Therefore, we hypothesized that EGL-1 may act cell-autonomously in the ALMR to promote exopher production. Given that high ectopic expression of *egl-1* induces apoptosis in wild-type animals and *ced-3* appears dispensable for exophers, we conducted cell-specific rescue of *egl-1* using *ced-3* as the genetic background. The *egl-1;ced-3* double mutant exhibited a significantly lower ALMR exopher rate compared to the *ced-3* single mutant (Figure 2C). We found that touch neuron-specific expression of EGL-1 with independent promoters (*Pmec-17::egl-1* and *Pmec-7::egl-1*) rescued the ALMR exopher rate in the *egl-1;ced-3* double mutant (Figure 2C). In summary, EGL-1 appears to act cell-autonomously in the ALMR neuron to promote exopher production, potentially through the mitochondrial protein CED-9, but not the downstream caspase CED-3.

### EGL-1 may spur exopher production via mitochondrial fission

The long-standing perception of *egl-1* as the BH3-only protein that is necessary and sufficient for triggering apoptosis was recently updated by a report of its non-apoptotic function in promoting mitochondrial fission in healthy cells. The studies conducted in embryos and muscles revealed that the EGL-1-CED-9 complex facilitates the recruitment of DRP-1 to mitochondria and promotes fission; while in the absence of EGL-1, CED-9 promotes FZO-1-dependent mitochondrial fusion^[21,22]^. Because endogenous EGL-1 has been found to function in the ALM neurons^[21]^, we speculated that *egl-1* also regulates mitochondrial dynamics in this neuron. We examined the mitochondrial morphology in the ALMR soma with a mitochondria-GFP reporter and categorized the ALMR soma mitochondrial morphology into three types: fragmented, tubular, or highly globular. Both *egl-1* and *ced-9* gain-of-function mutants exhibited more highly globular (fused), and less fragmented (fissioned) mitochondria compared to wild type (Figure 3A). These findings suggest that the EGL-1-CED-9 complex promotes mitochondrial fission in the ALMR neuron.

**Figure 3.**
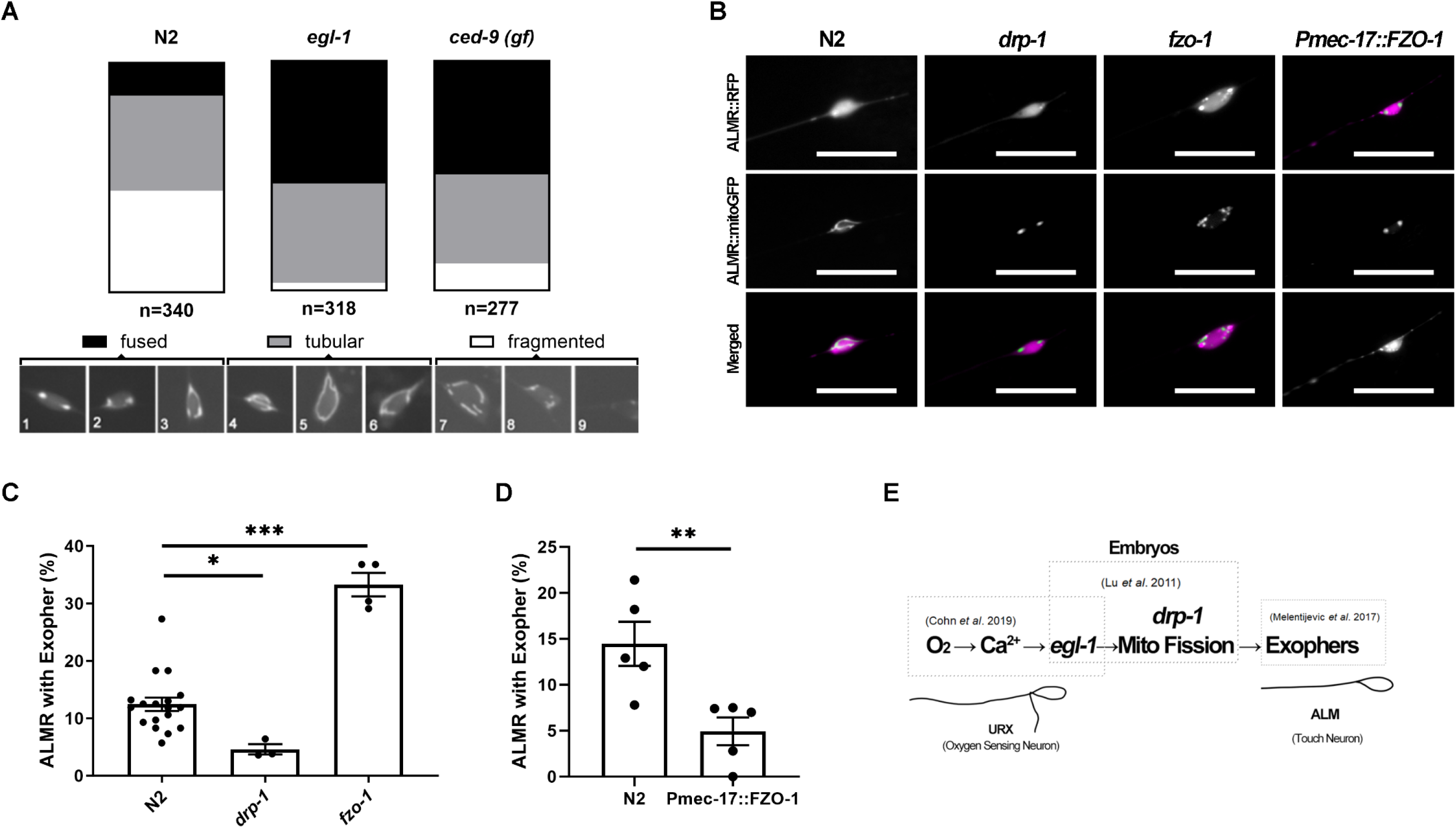
*egl-1*-regulated mitochondrial fission promotes exopher production in ALMR neuron. A. Mitochondria morphology is scored in ALMR soma in wild-type background, *egl-1lf(n1084n3082)* and *ced-9 950)* mutant animals. Both *egl-1* loss-of-function and *ced-9* gain-of-function mutants have more fused and ragmented mitochondria. All strains have the Is[Pmec-7::RFP];Is[Pmec-7::mitoGFP]. B. Representative image of mitochondria morphology in wild-type N2, *drp-1(tm1108), fzo-1(tm1133)* and the 1 overexpression strain *Ex[Pmec-17::FZO-1*Most of the N2 display tubular and circular mitochondria in ALMR. Both *drp-1* loss-of-function mutants and animals with touch-neuron-specific over-expression of FZO-1 y large, fused mitochondria aggregates in ALMR. *fzo-1* loss-of-function mutants exhibit fragmented hondria in ALMR. ]. All strains have the Is[Pmec-7::RFP];Is[Pmec-7::mitoGFP]. C. ALMR exophers are scored in *drp-1(tm1108), fzo-1(tm1133)* and compared to wild-type N2. *drp-1* loss-of-on mutants have less exophers, while fzo-1 loss-of-function mutants have more exophers. All strains the Is[Pmec-17::GFP]. Dots represent groups with N = 1082, 231 and 299 worms total for wild-type and nts by order, respectively. D. ALMR exophers are scored in wild-type N2 and the FZO-1 overexpression strain *Ex[Pmec-17::FZO-1]*. Cell-fic over-expression of FZO-1 suppressed the exopher production in ALMR. N = 311 and 184 for wild-type verexpression strain respectively. E. Summary showing linkage of three disparate studies for activity-dependent expression of *egl-1*, EGL-1 role ochondrial dynamics, and exophers. All animals are adult day 1. Scale bars: 20 μm.

Next, we wanted to investigate the relationship between mitochondrial fission and exopher production. As mentioned above, recent findings showed that mechanical force generated by the fertilized eggs in the uterus stimulates production of exopher in the touch neurons, so studying exophers in animals with excessive egg retention is problematic^[19]^. Considering that the mitochondrial fission and fusion mutants have reduced brood size and abnormally high rate of unfertilized eggs or egg retention, we scored a subset of worms with no excessive egg retention (day 1 adults with 10-20 fertilized eggs) for ALMR exophers (see methods). In the fission mutant *drp-1*, mitochondria appeared fused into highly globular puncta (Figure 3B) and showed a significantly lower ALMR exopher rate (Figure 3C). Conversely, the fusion mutant *fzo-1* displayed fragmented mitochondria (Figure 3B) and had a significantly higher ALMR exopher rate (Figure 3C). These findings suggest a correlation between mitochondrial fission and exopher production.

To test if the mitochondrial dynamics regulate exopher production cell-autonomously, we overexpressed the mitochondrial fusion protein FZO-1 in the touch neurons. We found that overexpressing FZO-1 in the touch neurons caused mitochondria to fuse into highly globular puncta (Figure 3B), and significantly decreased the ALMR exopher rate (Figure 3D). We conclude that the EGL-1 promotes exopher production in ALMR possibly by regulating mitochondrial fission-fusion dynamics through CED-9.

Despite the drastic morphological change and loss of cell content, exophers have been surprisingly found to be neuroprotective for the touch neurons^[9]^. Knowing that *egl-1* promotes exopher production and regulates mitochondrial dynamics in neurons, we next investigated the role of the endogenously expressed *egl-1* in URX neuronal health. We examined the URX dendrite and quantified the dendritic beading phenotype, a hallmark for compromised neuronal health in *C. elegans* studies^[23]^. When *egl-1* was knocked down selectively in URX, we found an increase in URX dendritic beading, which was suppressed by growing the animals in a 10% O_2_ environment (Figure 4A and C). These results suggest that the activity-dependent expression of *egl-1* confers beneficial effects on dendritic health when the URX neuron is chronically activated with high Ca^2+^ level.

**Figure 4.**
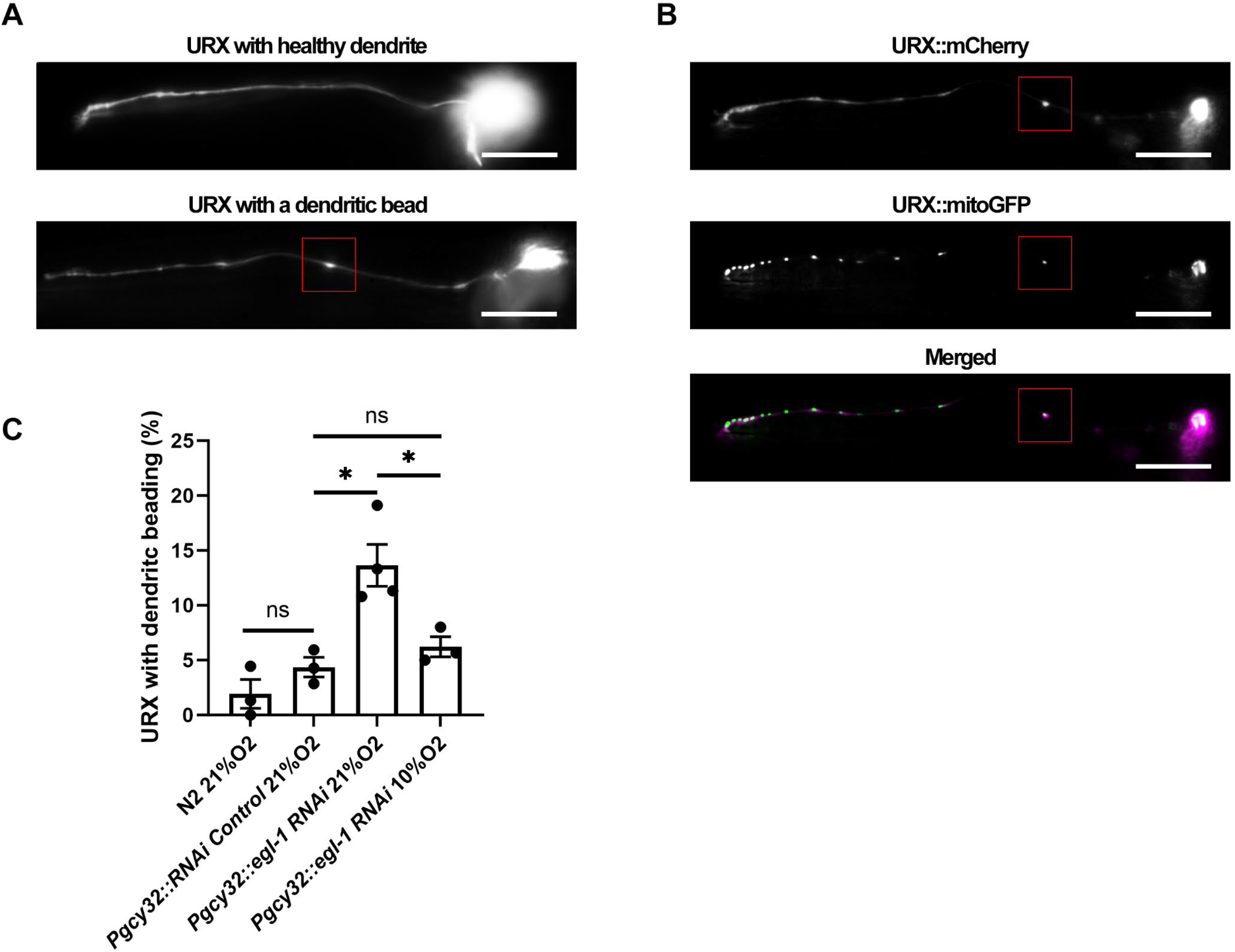
*egl-1* deficiency leads to dendritic beading in URX neuron. A. Representative image of URX with a healthy dendrite (top) and URX with dendritic beading (bottom). B. URX dendritic beads contain mitoGFP-labeled mitochondria (red box). URX and mitochondria are labeled *s[Pegl-1::mCherry]* and *Is[Pegl-1::mitoGFP]* reporters. C. URX dendritic beading is scored in wild-type N2,RNAi knockdown strains *Ex[Pgcy32::GFP, 32::mCherry RNAi]* and *Ex[Pgcy32::GFP, Pgcy32::egl-1 RNAi]*. URX-specific knockdown of *egl-1* ased dendritic beading, while the control strain with URX-specific control knockdown of mCherry appeared no different from the N2. Growth in 10% O_2_ environment suppressed the dendritic beading in the specific *egl-1* knockdown strain. Dots represent groups with N = 211, 233, 259 and 163 worms total for ype, RNAi control, *egl-1* RNAi, and *egl-1* RNAi in 10% O_2_ environment, respectively. Animals are adult day 1. Scale bars: 20 μm.

Mitochondria have been previously found to localize in the dendritic beads of touch neurons^[23]^. Interestingly, we also found mitochondria localization in the URX dendrite beads (Figure 4B). These results suggest that endogenous *egl-1* expression in URX is beneficial for the neuron, possibly through its mitochondrial function.

Valproic acid (VPA) is a frequently prescribed antiepileptic drug with application in various neurological and psychiatric disorders^[24,25]^. VPA is also reported to provide neuroprotective effects in mammalian studies^[26,27,28,29]^. In *C. elegans*, VPA was also found to extend lifespan and to protect against neurodegeneration^[30,31]^. We serendipitously found that VPA increases exopher production in multiple neurons, including URX, ALM, HSN and CEP (Supplemental Figure 3A,B and C). In addition, we found evidence that the VPA-induction of exophers in URX can be suppressed by reducing *egl-1* level. Both a low (10%) oxygen environment (where URX has low activity and low *egl-1* level) and URX-specific RNAi knock down of *egl-1*, suppressed the increase of exophers with VPA (Supplemental Figure 3D and E). These results indicate possible interactions among VPA, *egl-1*, and exopher production in maintaining neuronal health.

## Discussion

We found a new non-apoptotic role for the BH3-only protein EGL-1 where it promotes exopher production cell-autonomously and neuronal health, potentially through the regulation of mitochondrial dynamics. Our findings link three previously disparate studies on activity-dependent expression of *egl-1*, a role for *egl-1* in mitochondrial fission in embryo and muscle, and exophers (Figure 3E).

Exopher studies in *C. elegans* have revealed cellular-molecular mechanisms in exopher production and function. Various stresses, including fasting and oxidative stress, have been found to increase exopher production through multiple cell-nonautonomous signaling pathways^[32]^. Intermediate filaments that associate with aggresomes are required cell-autonomously for exopher production^[33]^. Additionally, the autophagy protein ATG-16.2 has been found to promote exopher production cell autonomously^[34]^. Complementing these findings in stress-response pathways, we discovered that the apoptosis trigger in *C. elegans*, the BH3-only protein EGL-1, has a non-apoptotic function in promoting exopher production cell-autonomously.

Intriguingly, exopher production appears to begin and end using cellular and molecular processes shared with apoptosis, while the middle caspase step appears dispensable. Exopher production, distinct from apoptosis, does not induce cell death in the non-stressed physiological conditions^[9]^. Consistent with previous findings, we found the conserved caspase CED-3, which executes apoptosis, is dispensable for exopher production. In some ways, exophers resemble apoptotic bodies, blebs that protrude and are released from apoptotic cells. However, exophers lack a level of phosphatidylserine (PS) on the membrane surface that is detectable by annexin V in apoptotic bodies. The phagocytosis pathway that engulfs the exopher and promotes exopher production appears to share parallel roles in the degradation and stimulation of apoptotic cells. This suggests involvement of a sub-apoptotic level of PS exposure^[9,12]^. PS exposure in apoptosis requires the cleavage of the scramblase CED-8 by the activated caspase CED-3^[35,36]^. The mitochondrial factor WAH-1 also promotes PS exposure through the scramblase SCRM-1 in apoptosis^[37]^. In apoptosis, WAH-1 is released from mitochondria in a CED-3-dependent manner; however, WAH-1 can be partially released in the CED-3 mutant under EGL-1 expression ^[38]^. Together in context with our findings, these results suggest that under non-apoptotic conditions, EGL-1 may promote PS exposure by facilitating mitochondrial release of WAH-1, thereby promoting exopher production.

BH3-only proteins facilitate the release of mitochondrial factors through mitochondrial outer membrane permeabilization (MOMP)^[39]^. MOMP, often studied in apoptosis, can also happen in non-apoptotic contexts. Several studies have revealed “minority MOMP”, where MOMP is limited to sublethal levels and is involved in cellular senescence^[40,41]^. Interestingly, DRP-1-mediated mitochondria fission facilitates both apoptotic MOMP and non-apoptotic minority MOMP^[42,43]^.Taken together, these studies and our results suggest that mitochondria dynamics may act downstream and/or in parallel with the BH3-only proteins to promote exopher PS exposure by promoting mitochondria-factor release through minority MOMP.

In addition to MOMP, BH3-only proteins and mitochondria dynamics are also found to function synergistically in other stress-induced pathways, such as calcium overload, mitochondrial permeability transition pore, and reactive oxygen species^[44,45,46,47]^. A recent study reported that mechanical force induces DRP1-mediated mitochondrial fission in cultured cells^[48]^. This discovery, in conjunction with our finding, fits with an independent recent *C. elegans* study that reported that mechanical force potentiates exopher extrusion^[19]^. In conclusion, EGL-1 and downstream mitochondrial dynamics in canonical apoptosis pathways have non-apoptotic roles in stress response pathways and exopher production.

Apoptosis is characterized by hallmarks such as membrane blebbing, nuclear condensation and fragmentation, mitochondrial fragmentation, PS exposure, and caspase activation^[49]^. Canonical apoptosis has been considered as a non-reversible cell-death pathway; however, recent studies have shown that cells can recover even after displaying apoptosis hallmarks^[50,51,52]^. Neurons can recover from stress-induced apoptosis even after displaying hallmarks including cell blebbing, mitochondria fragmentation and PS exposure, but cannot recover after caspase activation^[52]^. Given that exopher production in *C. elegans* is not dependent on the caspase CED-3, it will be interesting to investigate the relationship between exopher production and apoptosis pathways upstream of caspase activation in future studies.

We found that the exopher-promoting levels of endogenous *egl-1* appears beneficial for dendritic health, and the neuroprotective drug VPA promotes exopher production in multiple neurons. These findings are consistent with the previous research suggesting the benefits of exophers for neurons^[9]^. In a *C. elegans* Parkinson’s model study, knocking down *egl-1* enhanced dopaminergic neuron degeneration in aging animals^[53]^. Conversely, transgenic overexpression of *egl-1* in URX neurons induced cell death^[54]^. Therefore, both ectopically expressing high (lethal)-level *egl-1* and experimentally inhibiting the endogenously expressed low (nonlethal)-level *egl-1* are detrimental for neuronal health. We propose that under low to mild stress conditions, such as prolonged neuron activation and physiological aging, low *egl-1* expression helps to maintain neuronal health. We suspect that this may explain why tonically active sensory neurons (URX, AWC) in *C. elegans* endogenously express *egl-1* at surprising moderate levels without killing them^[8]^. Under high stress levels, high *egl-1* expression induces neuronal apoptosis, which is nevertheless relatively more controlled and less inflammatory comparing to other cell death pathways such as necrosis and ferroptosis. The canonical apoptosis hallmarks should be examined more carefully to distinguish ongoing apoptosis from stress-response mechanisms that have overlapping molecular pathways and cellular hallmarks with apoptosis.

Taking a broader view, because many therapeutic approaches for neurodegeneration target apoptosis to prevent cell death^[55]^, our study raises the possibility that blocking this early stage of apoptotic signaling may counterintuitively reduce the health of neurons. Additionally, exophers were also found to be important for the mitochondrial homeostasis cardiomyocytes^[56]^. Thus, these findings may have implications for fields of neurodegeneration, cancer, and heart disease.

## Materials and methods

### Animals

*C. elegans* strains were cultured at 21 °C on Nematode Growth Medium (NGM) plates and fed OP50 bacteria as described^[57]^. N2 was used as wild type. Additional strains are listed below. All genotypes were confirmed by PCR or sequencing.

JPS602 *vxEx602[Pgcy-32::GFP]*

JPS1809 *vxIs601[Pegl-1::mCherry::egl-1UTR];bcIs49[Pegl-1::mitoGFP]*.

JPS1031 *vxEx1031[Pgcy-32::mCherry]*

JPS1020 *egl-4(n477) IV; muIs102[Pgcy-32::GFP]*

JPS1148 *muIs102[Pgcy-32::GFP]*

JPS1810 *vxEx1512[Pgcy-32::GFP::unc-54UTR + Punc-122::RFP::unc-54UTR]*

JPS1806 *vx1513Ex[Pgcy-32::GFP, Pgcy-32::egl-1RNAi, Pmyo-3::mCherry]*

JPS1807 *vx1514Ex[Pgcy-32::GFP, Pgcy-32::mCherry RNAi, Pmyo-3::mCherry]*

TU2769 *uIs31[Pmec-17::gfp]*

JPS1712 *uIs31[Pmec-17::gfp] III; egl-1(n1084n3082) V*

JPS1713 *uIs31[Pmec-17::gfp] III; ced-3(ok2734) IV*

JPS1771 *uIs31[Pmec-17::gfp] III; ced-9(n1950) III*

JPS1714 *uIs31[Pmec-17::gfp] III; ced-3(ok2734) IV; egl-1(n1084n3082) V*

JPS1715 *uIs31[Pmec-17::gfp] III; ced-3(ok2734) IV; egl-1(n1084n3082) V; vxEx1510[Pmec-7::EGL-1, Pmyo-3::mCherry]*

JPS1716 *uIs31[Pmec-17::gfp] III; ced-3(ok2734) IV; egl-1(n1084n3082) V; vxEx1511[Pmec-17::EGL-1, Pmyo-3::mCherry]*

NM4244 *jsIs973 [Pmec-7::mRFP] III; jsIs609 [Pmec-7::mtGFP] X*

JPS1729 *jsIs973 [Pmec-7::mRFP] III; egl-1(n1084n3082) V; jsIs609 [Pmec-7::mtGFP] X*

JPS1770 *jsIs973 [Pmec-7::mRFP] III; ced-9(n1950) III; jsIs609 [Pmec-7::mtGFP] X*

JPS1731 *jsIs973 [Pmec-7::mRFP] III; drp-1(tm1108) IV; jsIs609 [Pmec-7::mtGFP] X*

JPS1732 *fzo-1(tm1133) II; jsIs973 [Pmec-7::mRFP] III; jsIs609 [Pmec-7::mtGFP] X*

JPS1727 *uIs31[Pmec-17::gfp] III; drp-1(tm1108) IV*

JPS1728 *fzo-1(tm1133) II; uIs31[Pmec-17::gfp] III*

JPS1739 *jsIs973 [Pmec-7::mRFP] III; jsIs609 [Pmec-7::mtGFP] X; vxEx1517[Pmec-17::FZO-1, Punc-122::GFP]*

BZ555 *egIs1[Pdat-1::GFP]*

JPS1666 *vsIs591[Ptph-1::GFP]*

### Strain Construction

Cell-specific RNAi knockdown strains were generated as described^[16]^. For cell-specific *egl-1* RNAi strain, an 876 bp *gcy-32* promoter was PCR fused with a 463 bp fragment of *egl-1* in the sense or antisense orientation. For control RNAi strain, the same promoter was PCR fused with a 528 bp fragment of mCherry from the plasmid pCFJ90, in the sense and antisense orientation. For both RNAi strains, constructs were transformed at 80 ng/μL each, with 5 ng/μL *Pmyo-3::mCherry* co-injection marker.

Cell-specific *egl-1* rescue strains were generated by PCR fusion. A 1366 bp *mec-7* promoter or a 1576 bp *mec-17* promoter was PCR fused with the *egl-1* gene (including the 3’UTR) amplified from N2 lysate. The constructs were injected at 20 ng/μL, with 5 ng/μL *Pmyo-3::mCherry* co-injection marker.

Cell-specific *fzo-1* overexpression strain was generated by PCR fusion. A 1576 bp *mec-17* promoter was PCR fused with the *fzo-1* gene (including the 3’UTR) amplified from N2 lysate. The construct was injected at 20 ng/μL, with 10 ng/μL *Punc-122::GFP* co-injection marker.

### Microscopy

Worms were immobilized with 10-30 mM sodium azide in NGM and mounted on 2% agarose pads for imaging. Epifluorescent images were captured on an Olympus IX51 inverted microscope equipped with an X-Cite FIRE LED Illuminator (Excelitas Technologies Corp.), using Olympus UPlanFL N 40X/0.75 NA, UPlanFL N 60X/1.25 Oil Ins objectives and QCapture Pro 6.0 software.

### Exopher Scoring

Exophers were scored in day 1 adults in a yes/no binary manner. ALMR exophers are scored according to the method in previous studies^[9,10]^. Animals with fewer than 10 or more than 20 fertilized eggs were excluded to minimize the non-specific effect of uterine occupation on ALMR exophers^[19]^. The presence of an URX exopher was scored when the soma exhibited an exopher budding (defined by size and shape) or a detached exopher(s) as shown in (Figure 1B). Both left and right URX neurons were inspected separately in each animal. The CEPD and HSN exophers were scored in the same manner as the URX exophers. Unpaired, planned t-tests were used in statistics. Data are presented in mean ± s.e.m.

### Mitochondria Morphology Scoring

ALMR Mitochondria were imaged with the *jsIs609 [Pmec-7::mtGFP]* tag and the photos were scored in blinded fashion. Mitochondria morphologies were scored as shown in (Figure 3A), morphologies represented by picture 1∼3, 4∼6, 7∼9 were categorized into globular, tubular, fragmented, respectively.

### Dendritic Beading Scoring

URX dendritic beading was scored in day 1 adults in a yes/no binary manner. Beading was scored when one or more bead-like GFP accumulation(s) was present (Figure 4A,B). The bead-like GFP accumulation was only counted when the bead diameter was thicker than adjacent dendritic segment. Both URXL and URXR were included.

### Oxygen Experiments

For 10% O_2_ experiments, animals were grown in a Modular Incubator Chamber (Billups-Rothenberg) attached to oxygen tanks containing 10% O_2_ balanced with nitrogen (Airgas).

### VPA treatment

Valproic acid (VPA, Sigma-Aldrich P4543) was added to the NGM plates. Fresh 9 mM VPA plates were prepared and used within a week.

## Acknowledgments

We thank the *Caenorhabditis* Genetics Center for strains. We appreciate the Pierce lab for advice, Susan Rozmiarek and Cory Gentry for technical assistance for feedback on the manuscript. We acknowledge receipt of reagents from Min Han. This work was funded in part by the NIH (R01GM122463, RF1AG057355, R21OD032463 - J.P.) Waggoner Fellowship for Alcohol Research (J.P.), David & Ellen Berman Fellowships for Huntington’s Research (J.P.), and F.M. Jones and H.L. Bruce Graduate Fellowship (Z.W.).

## Author contributions

Z.W. and J.P. designed research; Z.W. and E.A.C. performed research; Z.W. contributed new reagents/analytic tools; Z.W. analyzed data; J.P. and Z.W. acquired funding; and Z.W. and J.P. wrote the paper.

## References

1 Degterev, A., & Yuan, J. (2008). Expansion and evolution of cell death programmes. Nature reviews. Molecular cell biology, 9(5), 378–390. 10.1038/nrm2393

2 Conradt, B., & Horvitz, H. R. (1998). The C. elegans protein EGL-1 is required for programmed cell death and interacts with the Bcl-2-like protein CED-9. Cell, 93(4), 519–529. 10.1016/s0092-8674(00)81182-4

3 Conradt, B., & Horvitz, H. R. (1999). The TRA-1A sex determination protein of C. elegans regulates sexually dimorphic cell deaths by repressing the egl-1 cell death activator gene. Cell, 98(3), 317–327. 10.1016/s0092-8674(00)81961-3

4 Doerflinger, M., Glab, J. A., & Puthalakath, H. (2015). BH3-only proteins: a 20-year stock-take. The FEBS journal, 282(6), 1006–1016. 10.1111/febs.13190

5 O’Reilly, L. A., Cullen, L., Visvader, J., Lindeman, G. J., Print, C., Bath, M. L., Huang, D. C., & Strasser, A. (2000). The proapoptotic BH3-only protein bim is expressed in hematopoietic, epithelial, neuronal, and germ cells. The American journal of pathology, 157(2), 449–461. 10.1016/S0002-9440(10)64557-9

6 Lein, E. S., Hawrylycz, M. J., Ao, N., Ayres, M., Bensinger, A., Bernard, A., Boe, A. F., Boguski, M. S., Brockway, K. S., Byrnes, E. J., Chen, L., Chen, L., Chen, T. M., Chin, M. C., Chong, J., Crook, B. E., Czaplinska, A., Dang, C. N., Datta, S., Dee, N. R., … Jones, A. R. (2007). Genome-wide atlas of gene expression in the adult mouse brain. Nature, 445(7124), 168–176. 10.1038/nature05453

7 Hawrylycz, M. J., Lein, E. S., Guillozet-Bongaarts, A. L., Shen, E. H., Ng, L., Miller, J. A., van de Lagemaat, L. N., Smith, K. A., Ebbert, A., Riley, Z. L., Abajian, C., Beckmann, C. F., Bernard, A., Bertagnolli, D., Boe, A. F., Cartagena, P. M., Chakravarty, M. M., Chapin, M., Chong, J., Dalley, R. A., … Jones, A. R. (2012). An anatomically comprehensive atlas of the adult human brain transcriptome. Nature, 489(7416), 391–399. 10.1038/nature11405

8 Cohn, J., Dwivedi, V., Valperga, G., Zarate, N., de Bono, M., Horvitz, H. R., & Pierce, J. T. (2019). Activity-Dependent Regulation of the Proapoptotic BH3-Only Gene egl-1 in a Living Neuron Pair in Caenorhabditis elegans. G3 (Bethesda, Md.), 9(11), 3703–3714. 10.1534/g3.119.400654

9 Melentijevic, I., Toth, M. L., Arnold, M. L., Guasp, R. J., Harinath, G., Nguyen, K. C., Taub, D., Parker, J. A., Neri, C., Gabel, C. V., Hall, D. H., & Driscoll, M. (2017). C. elegans neurons jettison protein aggregates and mitochondria under neurotoxic stress. Nature, 542(7641), 367–371. 10.1038/nature21362

10 Arnold, M. L., Cooper, J., Grant, B. D., & Driscoll, M. (2020). Quantitative Approaches for Scoring in vivo Neuronal Aggregate and Organelle Extrusion in Large Exopher Vesicles in C. elegans. Journal of visualized experiments : JoVE, (163), 10.3791/61368. https://doi.org/10.3791/61368

11 van Niel, G., D’Angelo, G., & Raposo, G. (2018). Shedding light on the cell biology of extracellular vesicles. Nature reviews. Molecular cell biology, 19(4), 213–228. 10.1038/nrm.2017.125

12 Wang, Y., Arnold, M. L., Smart, A. J., Wang, G., Androwski, R. J., Morera, A., Nguyen, K. C. Q., Schweinsberg, P. J., Bai, G., Cooper, J., Hall, D. H., Driscoll, M., & Grant, B. D. (2023). Large vesicle extrusions from C. elegans neurons are consumed and stimulated by glial-like phagocytosis activity of the neighboring cell. eLife, 12, e82227. 10.7554/eLife.82227

13 Shinoda, H., Shannon, M., & Nagai, T. (2018). Fluorescent Proteins for Investigating Biological Events in Acidic Environments. International journal of molecular sciences, 19(6), 1548. 10.3390/ijms19061548

14 White, J. G., Southgate, E., Thomson, J. N., & Brenner, S. (1986). The structure of the nervous system of the nematode Caenorhabditis elegans. Philosophical transactions of the Royal Society of London. Series B, Biological sciences, 314(1165), 1–340. 10.1098/rstb.1986.0056

15 Busch, K. E., Laurent, P., Soltesz, Z., Murphy, R. J., Faivre, O., Hedwig, B., Thomas, M., Smith, H. L., & de Bono, M. (2012). Tonic signaling from O2 sensors sets neural circuit activity and behavioral state. Nature neuroscience, 15(4), 581–591. 10.1038/nn.3061

16 Esposito, G., Di Schiavi, E., Bergamasco, C., & Bazzicalupo, P. (2007). Efficient and cell specific knock-down of gene function in targeted C. elegans neurons. Gene, 395(1-2), 170–176. 10.1016/j.gene.2007.03.002

17 Hengartner, M. O., & Horvitz, H. R. (1994). C. elegans cell survival gene ced-9 encodes a functional homolog of the mammalian proto-oncogene bcl-2. Cell, 76(4), 665–676. 10.1016/0092-8674(94)90506-1

18 Yuan, J., Shaham, S., Ledoux, S., Ellis, H. M., & Horvitz, H. R. (1993). The C. elegans cell death gene ced-3 encodes a protein similar to mammalian interleukin-1 beta-converting enzyme. Cell, 75(4), 641–652. 10.1016/0092-8674(93)90485-9

19 Wang, G., Guasp, R., Salam, S., Chuang, E., Morera, A., Smart, A. J., Jimenez, D., Shekhar, S., Melentijevic, I., Nguyen, K. C., Hall, D. H., Grant, B. D., & Driscoll, M. (2024). Mechanical force of uterine occupation enables large vesicle extrusion from proteostressed maternal neurons. eLife, 13, RP95443. 10.7554/eLife.95443.1

20 Hammarlund, M., Hobert, O., Miller, D. M., 3rd, & Sestan, N. (2018). The CeNGEN Project: The Complete Gene Expression Map of an Entire Nervous System. Neuron, 99(3), 430–433. 10.1016/j.neuron.2018.07.042

21 Lu, Y., Rolland, S. G., & Conradt, B. (2011). A molecular switch that governs mitochondrial fusion and fission mediated by the BCL2-like protein CED-9 of Caenorhabditis elegans. Proceedings of the National Academy of Sciences of the United States of America, 108(41), E813–E822. 10.1073/pnas.1103218108

22 Tan, F. J., Husain, M., Manlandro, C. M., Koppenol, M., Fire, A. Z., & Hill, R. B. (2008). CED-9 and mitochondrial homeostasis in C. elegans muscle. Journal of cell science, 121(Pt 20), 3373–3382. 10.1242/jcs.032904

23 Toth, M. L., Melentijevic, I., Shah, L., Bhatia, A., Lu, K., Talwar, A., Naji, H., Ibanez-Ventoso, C., Ghose, P., Jevince, A., Xue, J., Herndon, L. A., Bhanot, G., Rongo, C., Hall, D. H., & Driscoll, M. (2012). Neurite sprouting and synapse deterioration in the aging Caenorhabditis elegans nervous system. The Journal of neuroscience : the official journal of the Society for Neuroscience, 32(26), 8778–8790. 10.1523/JNEUROSCI.1494-11.2012

24 Perucca E. (2002). Pharmacological and therapeutic properties of valproate: a summary after 35 years of clinical experience. CNS drugs, 16(10), 695–714. 10.2165/00023210-200216100-00004

25 Isoherranen, N., Yagen, B., & Bialer, M. (2003). New CNS-active drugs which are second-generation valproic acid: can they lead to the development of a magic bullet?. Current opinion in neurology, 16(2), 203–211. 10.1097/01.wco.0000063774.81810.30

26 Ren, M., Leng, Y., Jeong, M., Leeds, P. R., & Chuang, D. M. (2004). Valproic acid reduces brain damage induced by transient focal cerebral ischemia in rats: potential roles of histone deacetylase inhibition and heat shock protein induction. Journal of neurochemistry, 89(6), 1358–1367. 10.1111/j.1471-4159.2004.02406.x

27 Qing, H., He, G., Ly, P. T., Fox, C. J., Staufenbiel, M., Cai, F., Zhang, Z., Wei, S., Sun, X., Chen, C. H., Zhou, W., Wang, K., & Song, W. (2008). Valproic acid inhibits Abeta production, neuritic plaque formation, and behavioral deficits in Alzheimer’s disease mouse models. The Journal of experimental medicine, 205(12), 2781–2789. 10.1084/jem.20081588

28 Zádori, D., Geisz, A., Vámos, E., Vécsei, L., & Klivényi, P. (2009). Valproate ameliorates the survival and the motor performance in a transgenic mouse model of Huntington’s disease. Pharmacology, biochemistry, and behavior, 94(1), 148–153. 10.1016/j.pbb.2009.08.001

29 Monti, B., Gatta, V., Piretti, F., Raffaelli, S. S., Virgili, M., & Contestabile, A. (2010). Valproic acid is neuroprotective in the rotenone rat model of Parkinson’s disease: involvement of alpha-synuclein. Neurotoxicity research, 17(2), 130–141. 10.1007/s12640-009-9090-5

30 Evason, K., Collins, J. J., Huang, C., Hughes, S., & Kornfeld, K. (2008). Valproic acid extends Caenorhabditis elegans lifespan. Aging cell, 7(3), 305–317. 10.1111/j.1474-9726.2008.00375.x

31 Kautu, B. B., Carrasquilla, A., Hicks, M. L., Caldwell, K. A., & Caldwell, G. A. (2013). Valproic acid ameliorates C. elegans dopaminergic neurodegeneration with implications for ERK-MAPK signaling. Neuroscience letters, 541, 116–119. 10.1016/j.neulet.2013.02.026

32 Cooper, J. F., Guasp, R. J., Arnold, M. L., Grant, B. D., & Driscoll, M. (2021). Stress increases in exopher-mediated neuronal extrusion require lipid biosynthesis, FGF, and EGF RAS/MAPK signaling. Proceedings of the National Academy of Sciences of the United States of America, 118(36), e2101410118. 10.1073/pnas.2101410118

33 Arnold, M. L., Cooper, J., Androwski, R., Ardeshna, S., Melentijevic, I., Smart, J., Guasp, R. J., Nguyen, K. C. Q., Bai, G., Hall, D. H., Grant, B. D., & Driscoll, M. (2023). Intermediate filaments associate with aggresome-like structures in proteostressed C. elegans neurons and influence large vesicle extrusions as exophers. Nature communications, 14(1), 4450. 10.1038/s41467-023-39700-1

34 Yang, Y., Arnold, M. L., Lange, C. M., Sun, L. H., Broussalian, M., Doroodian, S., Ebata, H., Choy, E. H., Poon, K., Moreno, T. M., Singh, A., Driscoll, M., Kumsta, C., & Hansen, M. (2024). Autophagy protein ATG-16.2 and its WD40 domain mediate the beneficial effects of inhibiting early-acting autophagy genes in C. elegans neurons. Nature aging, 10.1038/s43587-023-00548-1. Advance online publication. https://doi.org/10.1038/s43587-023-00548-1

35 Suzuki, J., Denning, D. P., Imanishi, E., Horvitz, H. R., & Nagata, S. (2013). Xk-related protein 8 and CED-8 promote phosphatidylserine exposure in apoptotic cells. Science (New York, N.Y.), 341(6144), 403–406. 10.1126/science.1236758

36 Chen, Y. Z., Mapes, J., Lee, E. S., Skeen-Gaar, R. R., & Xue, D. (2013). Caspase-mediated activation of Caenorhabditis elegans CED-8 promotes apoptosis and phosphatidylserine externalization. Nature communications, 4, 2726. 10.1038/ncomms3726

37 Wang, X., Wang, J., Gengyo-Ando, K., Gu, L., Sun, C. L., Yang, C., Shi, Y., Kobayashi, T., Shi, Y., Mitani, S., Xie, X. S., & Xue, D. (2007). C. elegans mitochondrial factor WAH-1 promotes phosphatidylserine externalization in apoptotic cells through phospholipid scramblase SCRM-1. Nature cell biology, 9(5), 541–549. 10.1038/ncb1574

38 Wang, X., Yang, C., Chai, J., Shi, Y., & Xue, D. (2002). Mechanisms of AIF-mediated apoptotic DNA degradation in Caenorhabditis elegans. Science (New York, N.Y.), 298(5598), 1587–1592. 10.1126/science.1076194

39 Tait, S. W., & Green, D. R. (2010). Mitochondria and cell death: outer membrane permeabilization and beyond. Nature reviews. Molecular cell biology, 11(9), 621–632. 10.1038/nrm2952

40 Ichim, G., Lopez, J., Ahmed, S. U., Muthalagu, N., Giampazolias, E., Delgado, M. E., Haller, M., Riley, J. S., Mason, S. M., Athineos, D., Parsons, M. J., van de Kooij, B., Bouchier-Hayes, L., Chalmers, A. J., Rooswinkel, R. W., Oberst, A., Blyth, K., Rehm, M., Murphy, D. J., & Tait, S. W. G. (2015). Limited mitochondrial permeabilization causes DNA damage and genomic instability in the absence of cell death. Molecular cell, 57(5), 860–872. 10.1016/j.molcel.2015.01.018

41 Victorelli, S., Salmonowicz, H., Chapman, J., Martini, H., Vizioli, M. G., Riley, J. S., Cloix, C., Hall-Younger, E., Machado Espindola-Netto, J., Jurk, D., Lagnado, A. B., Sales Gomez, L., Farr, J. N., Saul, D., Reed, R., Kelly, G., Eppard, M., Greaves, L. C., Dou, Z., Pirius, N., … Passos, J. F. (2023). Apoptotic stress causes mtDNA release during senescence and drives the SASP. Nature, 622(7983), 627–636. 10.1038/s41586-023-06621-4

42 Brooks, C., Cho, S. G., Wang, C. Y., Yang, T., & Dong, Z. (2011). Fragmented mitochondria are sensitized to Bax insertion and activation during apoptosis. American journal of physiology. Cell physiology, 300(3), C447–C455. 10.1152/ajpcell.00402.2010

43 Cao, K., Riley, J. S., Heilig, R., Montes-Gómez, A. E., Vringer, E., Berthenet, K., Cloix, C., Elmasry, Y., Spiller, D. G., Ichim, G., Campbell, K. J., Gilmore, A. P., & Tait, S. W. G. (2022). Mitochondrial dynamics regulate genome stability via control of caspase-dependent DNA damage. Developmental cell, 57(10), 1211–1225.e6. 10.1016/j.devcel.2022.03.019

44 Geng, X., Harry, B. L., Zhou, Q., Skeen-Gaar, R. R., Ge, X., Lee, E. S., Mitani, S., & Xue, D. (2012). Hepatitis B virus X protein targets the Bcl-2 protein CED-9 to induce intracellular Ca2+ increase and cell death in Caenorhabditis elegans. Proceedings of the National Academy of Sciences of the United States of America, 109(45), 18465–18470. 10.1073/pnas.1204652109

45 Duan, C., Kuang, L., Hong, C., Xiang, X., Liu, J., Li, Q., Peng, X., Zhou, Y., Wang, H., Liu, L., & Li, T. (2021). Mitochondrial Drp1 recognizes and induces excessive mPTP opening after hypoxia through BAX-PiC and LRRK2-HK2. Cell death & disease, 12(11), 1050. 10.1038/s41419-021-04343-x

46 Kong, D., Xu, L., Yu, Y., Zhu, W., Andrews, D. W., Yoon, Y., & Kuo, T. H. (2005). Regulation of Ca2+-induced permeability transition by Bcl-2 is antagonized by Drpl and hFis1. Molecular and cellular biochemistry, 272(1-2), 187–199. 10.1007/s11010-005-7323-3

47 Fu, H., Zhou, H., Yu, X., Xu, J., Zhou, J., Meng, X., Zhao, J., Zhou, Y., Chisholm, A. D., & Xu, S. (2020). Wounding triggers MIRO-1 dependent mitochondrial fragmentation that accelerates epidermal wound closure through oxidative signaling. Nature communications, 11(1), 1050. 10.1038/s41467-020-14885-x

48 Helle, S. C. J., Feng, Q., Aebersold, M. J., Hirt, L., Grüter, R. R., Vahid, A., Sirianni, A., Mostowy, S., Snedeker, J. G., Šarić, A., Idema, T., Zambelli, T., & Kornmann, B. (2017). Mechanical force induces mitochondrial fission. eLife, 6, e30292. 10.7554/eLife.30292

49 Taylor, R. C., Cullen, S. P., & Martin, S. J. (2008). Apoptosis: controlled demolition at the cellular level. Nature reviews. Molecular cell biology, 9(3), 231–241. 10.1038/nrm2312

50 Geske, F. J., Lieberman, R., Strange, R., & Gerschenson, L. E. (2001). Early stages of p53-induced apoptosis are reversible. Cell death and differentiation, 8(2), 182–191. 10.1038/sj.cdd.4400786

51 Tang, H. L., Tang, H. M., Mak, K. H., Hu, S., Wang, S. S., Wong, K. M., Wong, C. S., Wu, H. Y., Law, H. T., Liu, K., Talbot, C. C., Jr, Lau, W. K., Montell, D. J., & Fung, M. C. (2012). Cell survival, DNA damage, and oncogenic transformation after a transient and reversible apoptotic response. Molecular biology of the cell, 23(12), 2240–2252. 10.1091/mbc.E11-11-0926

52 You, W., Zhou, T., Knoops, K., Berendschot, T. T. J. M., van Zandvoort, M. A. M. J., Germeraad, W. T. V., Benedikter, B., Webers, C. A. B., Reutelingsperger, C. P. M., & Gorgels, T. G. M. F. (2023). Stressed neuronal cells can recover from profound membrane blebbing, nuclear condensation and mitochondrial fragmentation, but not from cytochrome c release. Scientific reports, 13(1), 11045. 10.1038/s41598-023-38210-w

53 Kim, H., Perentis, R. J., Caldwell, G. A., & Caldwell, K. A. (2018). Gene-by-environment interactions that disrupt mitochondrial homeostasis cause neurodegeneration in C. elegans Parkinson’s models. Cell death & disease, 9(5), 555. 10.1038/s41419-018-0619-5

54 Chang, A. J., Chronis, N., Karow, D. S., Marletta, M. A., & Bargmann, C. I. (2006). A distributed chemosensory circuit for oxygen preference in C. elegans. PLoS biology, 4(9), e274. 10.1371/journal.pbio.0040274

55 Pemberton, J. M., Pogmore, J. P., & Andrews, D. W. (2021). Neuronal cell life, death, and axonal degeneration as regulated by the BCL-2 family proteins. Cell death and differentiation, 28(1), 108–122. 10.1038/s41418-020-00654-2

56 Nicolás-Ávila, J. A., Lechuga-Vieco, A. V., Esteban-Martínez, L., Sánchez-Díaz, M., Díaz-García, E., Santiago, D. J., Rubio-Ponce, A., Li, J. L., Balachander, A., Quintana, J. A., Martínez-de-Mena, R., Castejón-Vega, B., Pun-García, A., Través, P. G., Bonzón-Kulichenko, E., García-Marqués, F., Cussó, L., A-González, N., González-Guerra, A., Roche-Molina, M., … Hidalgo, A. (2020). A Network of Macrophages Supports Mitochondrial Homeostasis in the Heart. Cell, 183(1), 94–109.e23. 10.1016/j.cell.2020.08.031

57 Brenner S. (1974). The genetics of Caenorhabditis elegans. Genetics, 77(1), 71–94. 10.1093/genetics/77.1.71

