## Supplemental Info for "Non-apoptotic role of EGL-1 in exopher production and neuronal health in *Caenorhabditis elegans*"

**Supplemental Material**

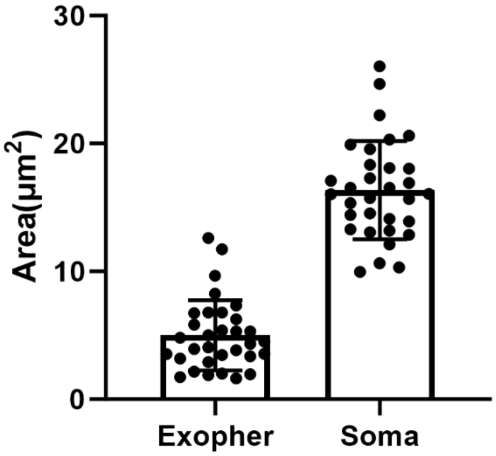


**Supplemental Figure 1.** Size of URX exophers and soma. Each dot represents measure from same neurons from independent worms. Data presented as mean ± SD.

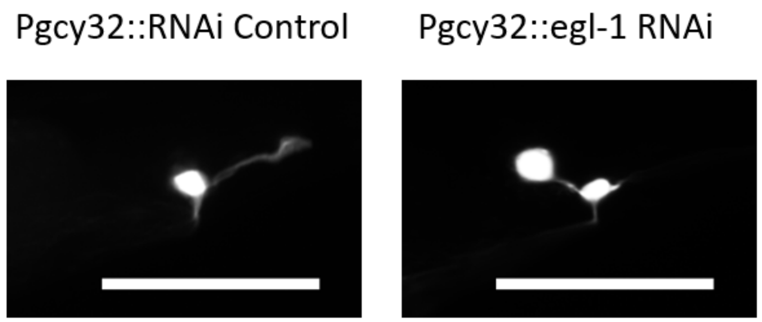


**Supplemental Figure 2.** Representative image of PQR neuron. The RNAi control strain (left) displays a PQR neuron with normal morphology. URX-specific *egl-1* knockdown strain (right) has an extra “undead” PQR soma. The suppression of apoptosis in PQR linage indicates the efficient knockdown of *egl-1* under the *gcy-32* promoter. PQR neurons labeled with *Ex[Pgcy32::GFP]* reporter.

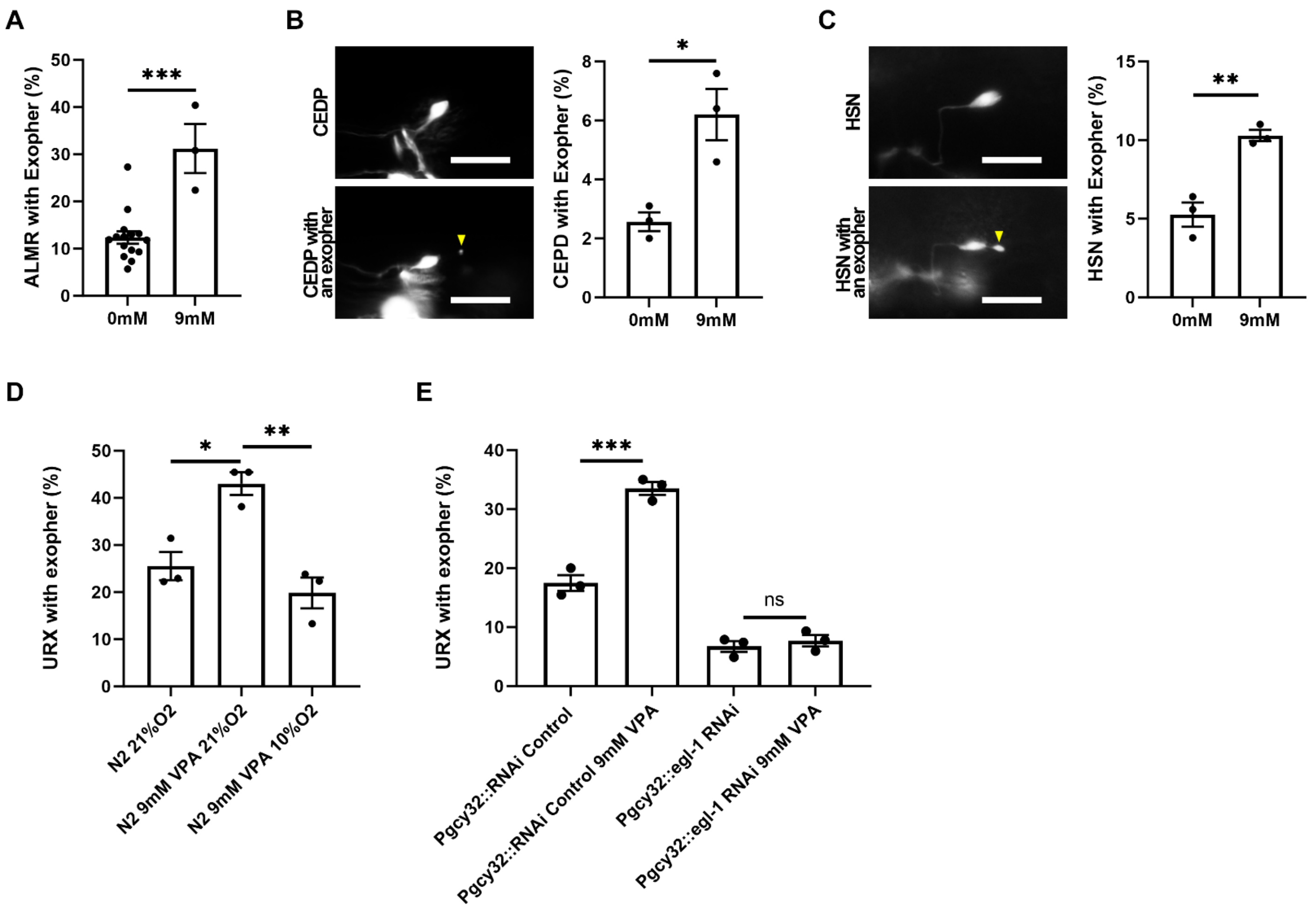


**Supplemental Figure 3. Neuroprotective valproic acid promotes exopher production in multiple neurons**

A-C. Representative images and quantification showing that VPA treatment increased exopher production ALMR (A), CEPD (B), and HSN (C) neurons compared to untreated controls. ALMR was visualized with *Is[Pmec-17::GFP],* CEP with *egIs1[Pdat-1::GFP]*, and HSN with *vsIs591[Ptph-1::GFP]*. N = 1082, 226 for control and VPA treatment in ALMR respectively. Dots represent groups with N = 263, 264 worms total for control and VPA treatment in CEPD respectively. N = 281, 488 for control and VPA treatment in HSN respectively.

D. VPA treatment increased URX exopher production. The VPA-increased exophers were suppressed when animals are grown in 10% $O_2$ environment, where URX has lower $Ca^{2+}$ activity and lower *egl-1* expression. URX visualized with *Ex[Pgcy32::GFP].* N = 185, 197 and 214 for control, VPA treatment and VPA treatment in 10%$O_2$ respectively.

E. URX-specific RNAi knockdown of *egl-1*, but not RNAi mCherry control, prevented VPA-induced increase in exopher production in URX neuron. Strains are *Ex[Pgcy32::GFP, Pgcy32::mCherry RNAi]* and *Ex[Pgcy32::GFP, Pgcy32::egl-1 RNAi].* N = 189, 235, 191 and 260 for RNAi control, RNAi control with VPA treatment, *egl-1* RNAi and *egl-1* RNAi with VPA treatment in 10% $O_2$ respectively.

Animals are adult day 1. Scale bars: 10 $\mu$m.
